# Shared phylogeographic structuring and recurrent hybridization underlie the diversification of *Microhyla* frog complexes in the Indochinese biodiversity hotspot

**DOI:** 10.64898/2026.08.18.745293

**Authors:** Christophe Dufresnes, Alexei V Trofimets, Vladislav A Gorin, Parinya Pawangkhanant, Nikita S Kliukin, Dmitriy V Arkhipov, Sơn Xuan Lê, Mahmudul Hasan, Mohd Abdul Muin, AA Thasun Amarasinghe, Amir Hamidy, Jinmin Chen, Yunhe Wu, Sengvilay Lorphengsy, Sang Ngoc Nguyen, Haipeng Zhao, Jieqiong Jin, Robert Murphy, Tan Van Nguyen, Spartak N Litvinchuk, Zhiyong Yuan, Jing Che, Chatmongkon Suwannapoom, Nikolay A Poyarkov

**Author notes:** corresponding authors: Christophe Dufresnes; Jing Che; Chatmongkon Suwannapoom; Nikolay A. Poyarkov. equal contribution (Christophe Dufresnes, Alexei V Trofimets).

## Abstract

Comparative phylogeography provides a powerful framework to identify the historical processes shaping biodiversity hotspots by testing whether co-distributed species exhibit shared patterns of diversification. Southeast Asia harbors exceptional biodiversity, yet the extent to which common paleogeographic and climatic drivers have structured diversification across taxa remains poorly understood. Here, we investigated the evolutionary history of five widespread *Microhyla* species complexes distributed across the Indochinese Peninsula and adjacent regions using dense mitochondrial sampling (1,388 individuals) combined with genome-scale ddRAD sequencing (280 individuals). Across all complexes, both approaches recovered multiple lineages and remarkably congruent phylogeographic breaks and contact/intergradation zones associated with major Indochinese regions, including Myanmar, the Tenasserim-Malay Peninsula, southern Vietnam, and northern Vietnam-southern China, supporting the hypothesis that Indochina functions as a mosaic of stable biogeographic units. However, lineage divergence varied substantially among complexes, suggesting that common biogeographic drivers interacted with species-specific demographic histories. Phylogenomic analyses and cyto-nuclear discordance further revealed historical introgression in every complex, indicating that diversification involved both long-term allopatric isolation and reticulate evolution. These findings portray the Indochinese biodiversity hotspot as a dynamic evolutionary system where cycles of fragmentation and reconnection have repeatedly reshaped lineage boundaries. Moreover, the complex phylogeographic structure recovered exemplifies the urgent need for taxonomic revisions, for which genome-scale data provide an essential framework to validate mitochondrial hypotheses and assess admixture patterns for species delimitation. Finally, regions such as the southern Annamites and Tenasserim Hills emerged as recurrent hotspots of genetic diversity across independent lineages, highlighting their importance for conserving not only species/lineage richness but also the evolutionary processes and adaptive potential that sustain biodiversity.

## Introduction

Southeast Asia harbors some of the highest levels of terrestrial biodiversity on Earth (Gentry 1992; Slik et al. 2015; Morley 2018), with its core region, the Indochinese Peninsula, supporting exceptional species richness and endemism (de Bruyn et al. 2014; Morley 2018). Broad-scale biogeographic syntheses have long emphasized contrasts between species communities inhabiting Indochina and adjacent southeast Asian regions, such as Sundaland, the Indo-Burmese highlands and southeast China (Inger & Voris 2001; Woodruff 2010), and within the peninsula, additional subdivisions have been matched to ecoregions (Poyarkov et al. 2021). For instance, distinct assemblages of amphibians and reptiles are found in the Mekong basin, the Annamite mountains along the Vietnamese-Lao border, and the Malay Peninsula, and within known species, strong phylogeographic structure and cryptic lineage diversity are being increasingly reported (Inger & Voris 2001; Woodruff 2010; Bain & Hurley 2011; Poyarkov et al. 2021, 2023). These observations suggest that Indochina functions as a mosaic of biogeographic units where diversification and persistence have repeatedly occurred.

The evolutionary mechanisms of this mosaic, however, remain insufficiently resolved. Diversification across the peninsula has likely been influenced by the interaction of complex topography with long-term geological and climatic dynamics linked to the Himalayan orogenesis, shifts in monsoon regimes, and Pleistocene changes in rainforest cover and sea level affecting connectivity (Wang et al. 2005; Clift & Plumb 2008; Cannon et al. 2009; Corlett & Primack 2011; Hall 2013; Raes et al. 2014). Whether such external drivers have structured biodiversity in a predictable and recurrent manner across taxa, or whether lineage-specific ecological traits and demographic histories have produced more idiosyncratic patterns, remains to be established. Moreover, because most phylogeographic studies so far relied predominantly on mitochondrial markers, the contribution of hybridization to diversification dynamics remains largely unexplored. Comparative phylogeography based on multilocus genomic data can provide a powerful framework to fill these gaps, particularly when applied to widespread species complexes that share similar ecological requirements and dispersal capacities, and are therefore expected to respond comparably to past environmental change.

In this context, narrow-mouthed frogs of the genus *Microhyla* (Anura: Microhylidae) offer a promising model system. These small-bodied amphibians are widely distributed across South, Southeast, and East Asia, where they occupy a broad range of lowland and montane habitats (Gorin et al. 2020, 2021). Molecular surveys have revealed substantial mitochondrial diversity within several traditionally recognized widespread species, suggesting that some may in fact represent species complexes (Gorin et al. 2020). The genus has consequently undergone extensive taxonomic revision in recent years (e.g., Poyarkov et al. 2020; Hoang et al. 2021, 2022; Garg et al. 2022; Trofimets et al. 2024), and more than 50 *Microhyla* species are currently recognized (Frost 2026). While many species have restricted, often micro-endemic distributions, others share broad geographic ranges spanning multiple ecoregions and frequently occur in sympatry – up to seven *Microhyla* species may co-exist in some localities (NAP pers. obc.). These provide an ideal framework for comparative phylogeographic analyses aimed at assessing whether diversification within the Indochinese hotspot reflects shared historical processes or lineage-specific trajectories.

To address this question, we inferred phylogeographic structure and diversity in five *Microhyla* species or species complexes co-distributed across Southeast Asia, combining extensive mitochondrial sampling with multilocus analyses of genomic data. These clades branch at various positions in the *Microhyla* tree, therefore being broadly representative of the genus diversification within the last millions of years (Gorin et al. 2020). If their evolution was shaped by shared paleogeographic and climatic drivers, co-distributed lineages should exhibit congruent phylogeographic breaks, with their diversification unfolding in parallel. Alternatively, if species show independent or idiosyncratic phylogeographic patterns, this would indicate a stronger role of species-specific ecological preferences and historical contingencies in their diversification history.

## Methods

### Study system and sampling

We focused on five widespread *Microhyla* clades that diverged from one another approximately 20 to 30 million years ago (Ma) (Gorin et al. 2020): (i) *M. berdmorei* complex; (ii) *M. butleri* complex; (iii) *M. pulchra*; (iv) *M. fissipes* complex; and (v) *M. heymonsi* complex. With the exception of *M. pulchra*, these clades were suggested to harbor strong mitochondrial structure (> 5 Ma) consistent with the presence of cryptic species (Gorin et al. 2020), as recently demonstrated for the *M. berdmorei* complex (Trofimets et al. 2024).

To compare phylogeographic patterns of these groups, specimens were collected across their distributions during field expeditions conducted between 2007 and 2023 under collecting permits (Supplementary Table S1). DNA was extracted with the Qiagen DNeasy Blood and Tissue Kit and new genetic data were generated for 1,016 individuals. Combined with previously published data, our study focused on 1,420 different individuals.

### Mitochondrial analyses

We generated 1,325 mitochondrial DNA sequences from 947 specimens (937 new ones), including fragments of 12S ribosomal RNA gene (12S; up to 978 bp aligned), 16S ribosomal RNA gene (16S; up to 1,475 bp aligned, including adjacent tRNA), and cytochrome c oxidase subunit I gene (COI; up to 720 bp aligned) (Supplementary Table S2), following methods from Gorin et al. (2020). An additional 617 sequences from 451 individuals were retrieved from GenBank. In total, 1,942 sequences from 1,388 different individuals were analyzed for mtDNA. For each individual, sequences were concatenated and aligned in SeaView v5 (Gouy et al. 2021).

Analyses were conducted separately for each *Microhyla* clade, with sequences representative of lineages from other clades included as outgroups. The mitochondrial diversity was first explored by maximum-likelihood (ML) analyses in IQ-TREE v2.3.6 (Minh et al. 2020), using gene partitions (and codon position partitions for the coding COI), model finder and 1,000 ultrafast bootstraps (UFBs). We next evaluated tree topology and divergence times based on reduced datasets representing the major mitogroups identified (Supplementary Table S3). We performed Bayesian phylogenetic inferences in BEAST v2.7.7 (Bouckaert et al. 2019) with substitution models set to HKY+G, optimized relaxed clocks and a Yule tree prior. Time calibration was enforced by constraining the root age on the time to the most recent common ancestor (tMRCA) with normally distributed priors set to replicate the 95% posterior distributions reported for the corresponding nodes by Gorin et al. (2020), who inferred a genus-wide timetree from fossil and geological calibrations (Supplementary Table S4). Chains were run for 50M iterations, sampling every 5M, stationarity was monitored in Tracer v1.7.2 (Rambaut et al. 2018), maximum clade credibility trees (MCCTs) were produced with the *TreeAnnotator* BEAST module, discarding the first 10% of trees as burn-in.

### Genomic analyses

We generated genomic data using double-digest Restriction-site Associated DNA sequencing (ddRAD-seq) for 280 specimens (Supplementary Table S5), following the protocol described in Dufresnes et al. (2025). In a nutshell, library preparation involved restriction with *MseI* and *SbfI*, ligation of Illumina adapters with unique individual barcodes, PCR amplification, and size selection of 400–500 bp fragments. Samples were processed across four libraries sequenced paired-end (PE150) on a DNBSeq platform at BGI China, which yielded between 279M and 461M reads.

Bioinformatic processing was conducted using Stacks v2.60 (Catchen et al. 2013). Raw reads were demultiplexed with *process_radtags*, including adapter removal, quality filtering (Phred score > 30), and trimming to 100 bp. Loci were assembled and catalogued de novo using the *denovo.pl* pipeline with default parameters, which appear appropriate for downstream phylogenomic analyses (Rancilhac et al. 2023). Separate assemblies were made for each *Microhyla* clade (plus 2–4 outgroups for each). Final catalogs comprised 796,624–1,883,359 loci, with mean effective per-sample coverages of 18.1–25.9×. SNPs were called using the *populations* module, with the filtering parameter -p (the number of genotyped samples or predefined groups) and -r (the proportion of genotyped samples within a predefined group) adjusted for each dataset to minimize locus dropout and missing data. To control for putative PCR artefacts and overmerged paralogous loci, we respectively applied minimum allele frequency (-min-maf 0.05) and maximum heterozygosity thresholds (-max-obs-het 0.75), which however had no impact, suggesting little sensitivity to these potential issues. Outputs were obtained in various formats for downstream analyses (-vcf, - structure, -phylip-var-all) and were converted with custom R scripts when needed. The numbers of loci, SNPs and samples of each dataset are summarized in Table 1.

**Table 1.** Details of the datasets used for each analysis in each *Microhyla* species/species complexes. n: number of samples; og: outgroup(s); -p: minimum number of genotyped samples (*or predefined groups) per RAD locus; -r: minimum proportion of genotyped samples in each predefined group per RAD locus (when -p refers to predefined groups); -wss: -write-single-snp option (calling a single SNP per RAD locus); mtDNA phylogeny: ML reconstruction of mitochondrial sequences with IQ-TREE: mtDNA timetree: Bayesian time-calibrated reconstruction of representative mitochondrial sequences with BEAST; RAD phylogeny: ML reconstruction of RAD sequences with IQ-TREE; RAD clustering: Bayesian clustering with STRUCTURE; RAD H_o_: observed heterozygosity estimates with the Stacks *populations* module; RAD species tree/*f_b_*: multispecies coalescent and ABBA-BABA tests of selected RAD samples by SNAPPER and Dsuite.

|  | Analysis | Sample/parameter sets | datasets |
| --- | --- | --- | --- |
| *M. berdmorei* complex | mtDNA phylogeny | n = 126 + 16 og | 3,091 bp (12S+16S+COI) |
|  | mtDNA timetree | n = 10 + 1 og | 3,064 bp (12S+16S+COI) |
|  | RAD phylogeny | n = 36 + 3 og (-p 36) | 409,799 bp (1,896 loci, 8,170 SNPs) |
|  | RAD clustering | n = 36 (-p 36 -wss) | 1,297 SNPs |
| | RAD $H_o$ | n = 36 (-p 36) | 5,478 SNPs |
| | RAD species tree/$f_b$ | n = 17 + 3 og (-p 8 -r 0.5)* | 4,773 SNPs |
| *M. butleri* complex | mtDNA phylogeny | n = 286 + 16 og | 3,007 bp (12S+16S+COI) |
|  | mtDNA timetree | n = 12 + 1 og | 2,967 bp (12S+16S+COI) |
|  | RAD phylogeny | n = 50 + 2 og (-p 43) | 379,699 bp (1,705 loci, 9,741 SNPs) |
|  | RAD clustering | n = 50 (-p 43 -wss) | 1,311 SNPs |
| | RAD $H_o$ | n = 50 (-p 43) | 8,119 SNPs |
| | RAD species tree/$f_b$ | n = 19 + 2 og (-p 8 -r 0.5)* | 2,488 SNPs |
| *M. pulchra* | mtDNA phylogeny | n = 52 + 17 og | 851 bp (16S) |
|  | mtDNA timetree | n = 4 + 2 og | 851 bp (16S) |
|  | RAD phylogeny | n = 18 + 4 og (-p 18) | 981,471 bp (4,573 loci, 23,456 SNPs) |
|  | RAD clustering | n = 18 (-p 18 -wss) | 3,668 SNPs |
| | RAD $H_o$ | n = 18 (-p 18) | 19,214 SNPs |
| | RAD species tree/$f_b$ | n = 10 + 3 og (-p 5 -r 0.5)* | 4,295 SNPs |
| *M. fissipes* complex | mtDNA phylogeny | n = 292 + 9 og | 3,161 bp (12S+16S+COI) |
|  | mtDNA timetree | n = 14 + 1 og | 3,159 bp (12S+16S+COI) |
|  | RAD phylogeny | n = 82 + 3 og (-p 70) | 537,743 bp (2,310 loci, 3,987 SNPs) |
|  | RAD clustering | n = 82 (-p 70 -wss) | 1,480 SNPs |
| | RAD $H_o$ | n = 82 (-p 70) | 3,747 SNPs |
| | RAD species tree/$f_b$ | n = 15 + 3 og (-p 7 -r 0.5)* | 2,404 SNPs |
| *M. heymonsi* complex | mtDNA phylogeny | n = 632 + 17 og | 3,007 bp (12S+16S+COI) |
|  | mtDNA timetree | n = 15 + 1 og | 2,966 bp (12S+16S+COI) |
|  | RAD phylogeny | n = 92 + 4 og (-p 75) | 382,666 bp (1,744 loci, 10,800 SNPs) |
|  | RAD clustering | n = 92 (-p 75 -wss) | 1,514 SNPs |
| | RAD $H_o$ | n = 92 (-p 75) | 9,752 SNPs |
| | RAD species tree/$f_b$ | n = 23 + 3 og (-p 9 -r 0.5)* | 1,896 SNPs |

For each *Microhyla* clade, we investigated genetic structure, diversity and recent admixture based on all samples as follows. First, ML phylogenies were inferred in IQ-TREE from concatenated sequence alignments (including outgroup samples), using ModelFinder and 1,000 UFBs. Second, genetic structure among ingroup samples was inferred using STRUCTURE v2.3.4 (Pritchard et al. 2000), based on SNP datasets constrained to a single SNP per RAD locus (-write-single-snp flag). Analyses were performed under the admixture model with correlated allele frequencies among populations. To improve chain mixing and facilitate convergence at higher K values, the λ parameter was estimated and the Metropolis update frequency for Q was set to 1. Markov chains were run for 10,000 iterations following a burn-in of 10,000 iterations, which appeared sufficient to reach stationarity. Rather than relying on a statistically inferred “optimal” number of clusters (K), which is strongly influenced by sampling design, hierarchical structure, and the scale of inference, we explored multiple K values and focused on the highest K producing biologically interpretable and phylogeographically coherent clusters with respect to the phylogenomic analyses. For each K, multiple replicate runs were examined and evaluated based on log-likelihood values (ln(P|D)), admixture parameter estimates (α), pairwise F_st_ values among inferred clusters, and the representation of all clusters in individual ancestry proportions. Preferred clustering solutions were those maximizing ln(P|D) and inter-cluster differentiation while minimizing admixture and avoiding “ghost” clusters, i.e., clusters with negligible assignment probabilities. Third, observed heterozygosity (H_o_) was estimated directly by the *populations* module in Stacks when calling all informative SNPs, and computed from variable sites only.

We next conducted additional analyses to infer phylogenomic relationships, divergence times and historical gene flow, based on subsets of samples representative of the inferred nuclear lineages – selected based on high sequencing coverage and absence of admixture in clustering analyses (Supplementary Table S6). Rather than phylogenetic reconstructions based on concatenated alignments, which estimate average gene divergence times and absorb incomplete lineage sorting (ILS) and discordance among loci into branch lengths, we employed multispecies coalescent approaches that explicitly model gene-tree discordance and estimate species/lineage divergence times independently of underlying ancestral coalescent variation. Time-calibrated species trees were reconstructed with SNAPPER v1.1.5 (Stoltz et al. 2021), as implemented in BEAST v2.7.7. Input XML files were prepared in BEAUTi using a Yule tree prior, a tMRCA prior identical to that used for the mitochondrial timetrees (Supplementary Table S4), and gamma priors on the birth-rate (α = 2.0, β = 0.5) and coalescent-rate parameters (α = 2.0, β = 1.0). Because the scale parameter (β) of these priors can influence inferred coalescent depths and divergence times, we evaluated prior sensitivity in preliminary analyses using alternative prior combinations corresponding to broader or shallower coalescent histories and larger or smaller effective population sizes; these yielded qualitatively similar results. Chains were run for at least 1M iterations, sampling every 1,000, monitored in Tracer, and MCCTs were produced with *TreeAnnotator*, discarding the first 10% as burn-in. Finally, we assessed historical gene flow using *f*-statistics implemented in Dsuite v0.5 r58 (Malinsky et al. 2021). ABBA–BABA tests were performed with the *Dtrios* function across all trios (composed of two sister and one non-sister lineage), as defined by the inferred species tree topology. Significant excesses of shared derived alleles were subsequently summarized across the phylogeny using the *Fbranch* function and visualized as *f*-branch plots with the corresponding Python script distributed with the program.

## Results

### Phylogeographic structure, admixture and genetic diversity

Mitochondrial and genomic datasets recovered genetic structure across all five *Microhyla* clades, with recurrent phylogeographic breaks consistently separating major regions of Southeast Asia, including the southeastern China-Taiwan region, Indo-Burma, the northern and southern Tenasserim-Malay Peninsula, different sectors of the Annamite Mountains, and Sundaland (Figs. 1–2). Observed heterozygosity (H_o_) estimates, which reflect intra-populational genetic variation, showed marked geographic differences, with diversity hotspots frequently located in southern Vietnam and western Thailand, while coldspots are rather found in latitudinal extremes of the distributions (e.g., Taiwan, Myanmar, Sundaland region) (Fig. 3). H_o_ estimates are given in Table S5.

**Fig. 1.**
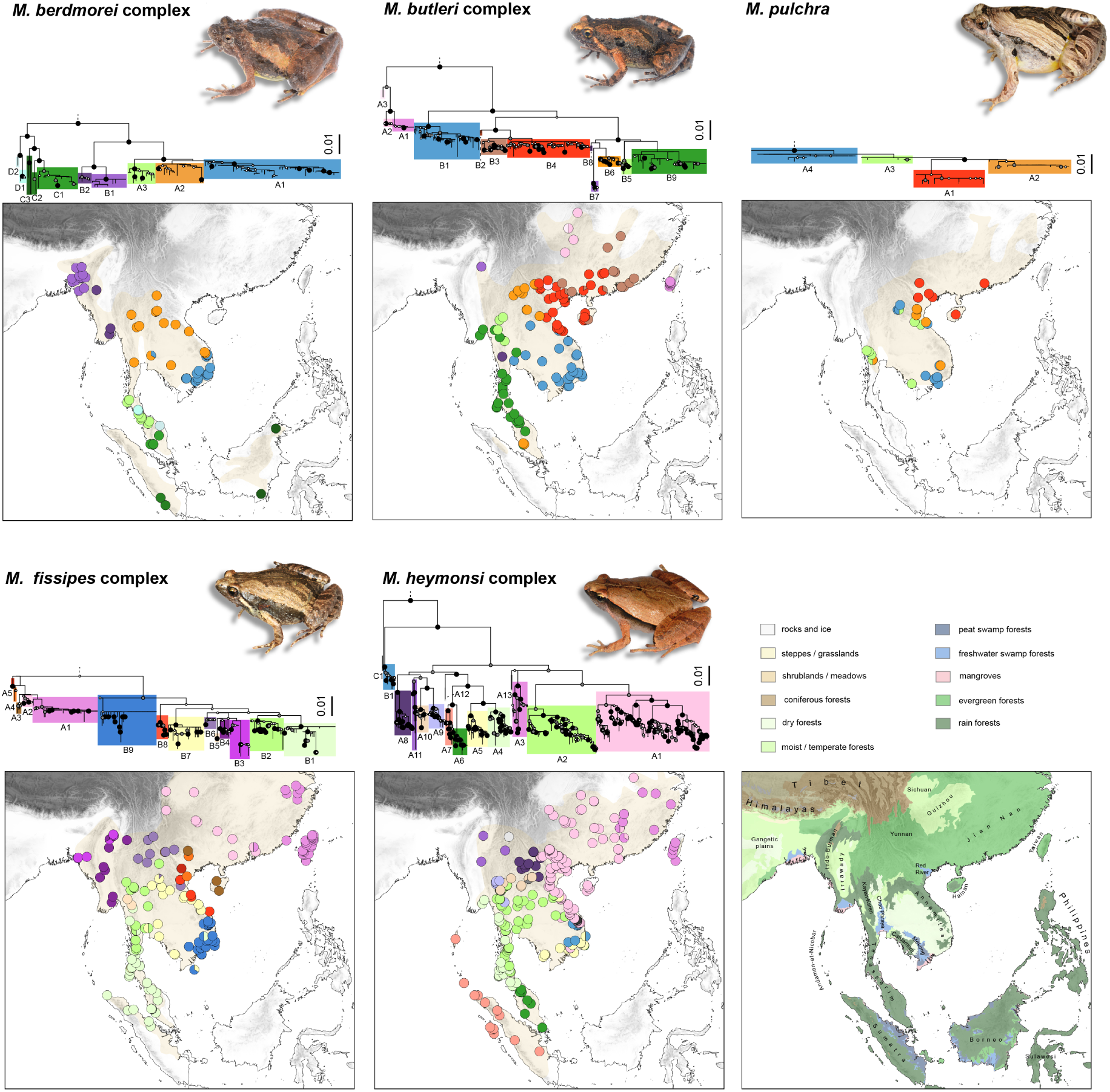
Mitochondrial phylogeography of *Microhyla* species/species complexes. Trees are ML reconstructions of 12S-16S-COI concatenated alignments; major mitogroups are highlighted in colors; node circles show branch support based on 1,000 ultrafast bootstraps (large black: ≥99; grey: 95-98; white: 70–94); the same scale is used for comparison (scale bars in substitutions/site); outgroups are not shown. Maps feature sampled localities, colored by major mitogroups; transparent yellow shades show species distributions. The bottom-right map shows major ecoregion types (Dinerstein et al. 2017) and geographic areas.

**Fig. 2.**
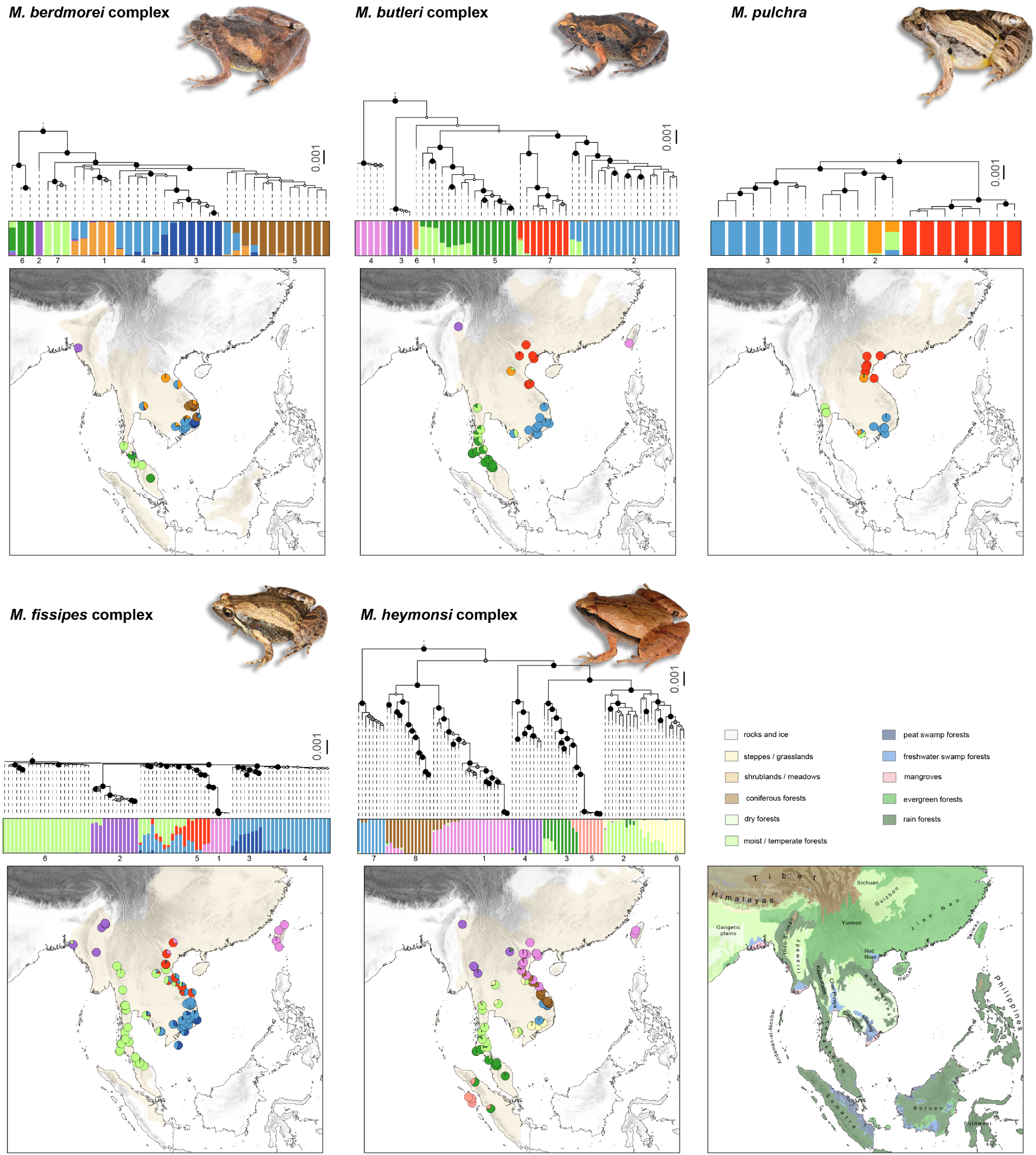
Genomic phylogeography of *Microhyla* species/species complexes. Trees are ML reconstructions of concatenated RAD loci; node circles show branch support based on 1,000 ultrafast bootstraps (large black: ≥99; grey: 95-98; white: 70-94); the same scale is used for comparison (scale bars in substitutions/site); outgroups are not shown. Barplots show ancestry to genetic clusters inferred by STRUCTURE, based on RAD SNPs. Maps show sampling localities colored by average ancestry; transparent yellow shades show species distributions. The bottom-right map shows major ecoregion types (Dinerstein et al. 2017) and geographic areas.

**Fig. 3.**
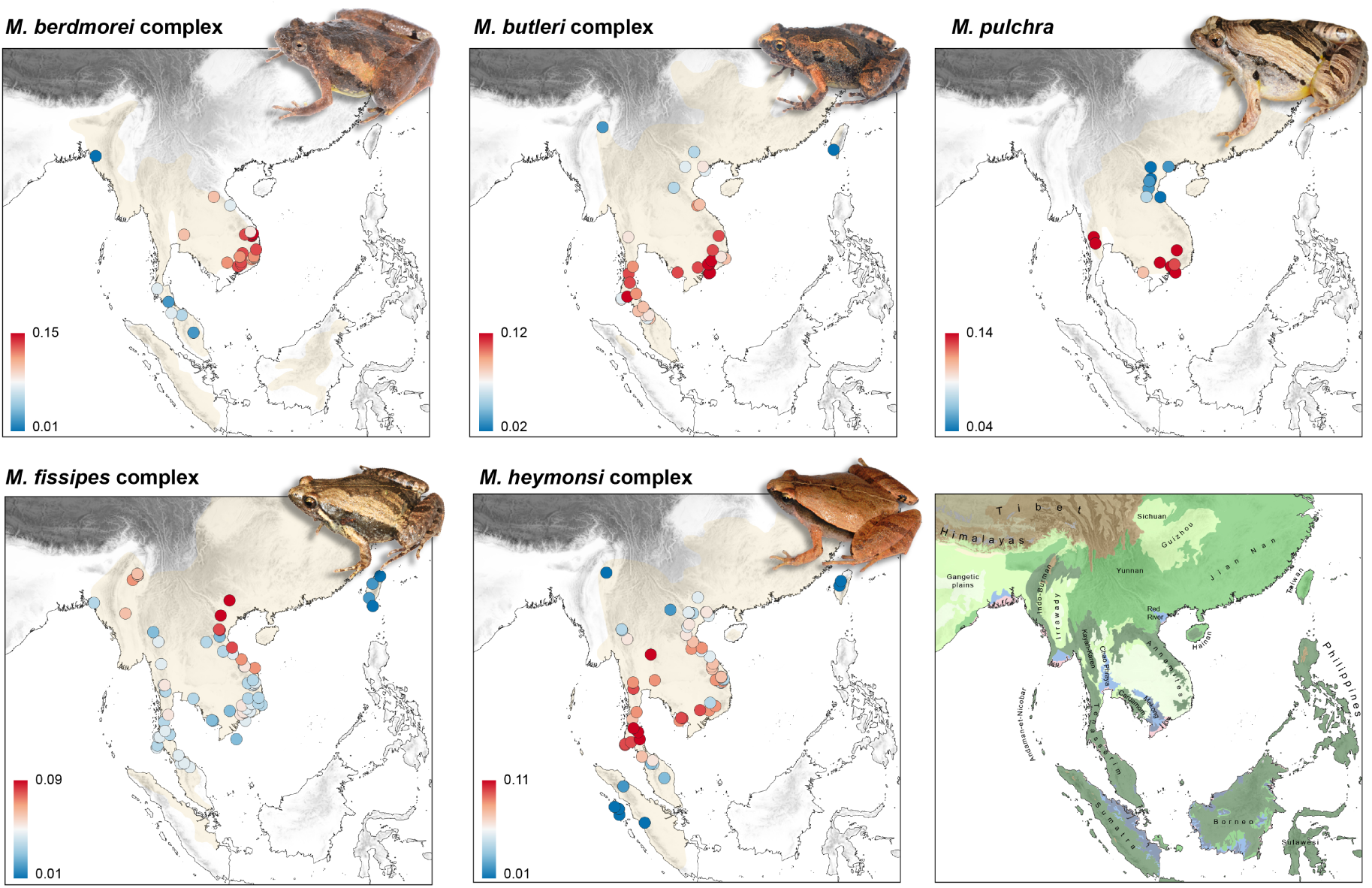
Genetic diversity in *Microhyla* species/species complexes. Sampling localities are colored by observed heterozygosity (H_o_). The bottom-right map shows major ecoregion types (Dinerstein et al. 2017) and geographic areas (see legend in Figs. 1–2).

The depth of diversification varied markedly among species complexes, ranging from shallow regional structuring in *M. pulchra* to extensive lineage fragmentation in the *M. heymonsi* complex. Moreover, although mitochondrial and genomic patterns were generally concordant, some lineages exhibited substantially weaker nuclear relative to mtDNA differentiation, most notably in the *M. fissipes* complex. In contrast, some deep nuclear lineages showed comparatively little mitochondrial divergence, such as the Myanmar lineage of *M. butleri*.

Across all clades, widespread genetic admixture and shifted mitochondrial compared to nuclear transitions indicate secondary contact and introgressive hybridization. These contact zones were often broadly located in the central Indochinese region – spanning the dry forests in eastern Thailand and Cambodia, the Cardamom Mountains, and the Mekong and Chao Phraya basins – as well as the northern Annamites and the central Tenasserim range.

Within each *Microhyla* clade, phylogeographic patterns can be summarized as follows.

In the *M. berdmorei* complex (mtDNA: n = 126; genomics: n = 36), four deeply divergent mitogroups corresponding to recently delimited species were consistently supported by genomic data (see also Trofimets et al. 2024) in western Indochina (*M. malcolmi*), Indo-Burma (*M. berdmorei*), the Malay Peninsula (*M. peninsularis*) and Sundaland (*M. sundaica*), with additional regional structuring and widespread admixture within each lineage, particularly in southwestern *M. malcolmi*. Observed heterozygosity broadly followed a longitudinal gradient, peaking in southern Vietnam and declining westward toward Myanmar and the Malay Peninsula.

In the *M. butleri* complex (mtDNA: n = 286; genomics: n = 50), both datasets supported a major phylogeographic split between Taiwan/China and Indochina. Indochinese populations further showed strong regional structuring across southeastern Indochina, the northern Annamites and nearby China (Hainan), Myanmar, Laos, the Kayah-Karen range, and the Malay Peninsula, with broad concordance between mitochondrial and genomic spatial patterns. Admixture zones are found in the Tenasserim and Cardamom ranges, and H_o_ shows a latitudinal pattern, with southern hotspots (southern Vietnam, Malay Peninsula) and northern coldspots (Myanmar, northern Vietnam, Taiwan).

*Microhyla pulchra* (mtDNA: n = 52; genomics: n = 18) exhibited shallow phylogeographic structure. Both datasets recovered weak but geographically coherent differentiation among northern Vietnam/southern China, the Annamites, western Thailand and Laos, with partial correspondence between mitochondrial and genomic groups. Admixture was detected in the Cardamom Mountains, and H_o_ contrasted between genetically diverse southern populations and genetically depauperate northern populations.

The *M. fissipes* complex (mtDNA: n = 292; genomics: n = 82) displayed strong mitochondrial diversification but shallow nuclear structure. MtDNA recovered two major mitogroups corresponding to *M. fissipes* and *M. mukhlesuri*, subdivided into regional lineages spanning Taiwan and mainland China, Myanmar, western Thailand, Sundaland and the Annamites, whereas genomic analyses resolved only weak geographic differentiation. Extensive admixture among genomic clusters was detected, particularly along the Annamite range and in southwestern China. In contrast to the other complexes, H_o_ peaked in northern Vietnam and remained relatively high in Myanmar, while being comparatively low elsewhere.

The *M. heymonsi* complex showed the strongest phylogeographic structure (mtDNA: n = 632; genomics: n = 92), with several lineages corresponding to recently described species (Hoang et al. 2021, 2022; Garg et al. 2022). Southern Vietnam harbors micro-endemic lineages/taxa, including *M. xodangorum* (C1), *M. daklakensis* (B1) and *M. ninhthuanensis* (A5). Additional lineages, generally supported by genomic analyses, were associated with the Annamites, Myanmar, western Thailand and Tenasserim, Peninsular Malaysia, Sundaland, northern Laos/Vietnam and southern China/Taiwan, including *M. nakkavaram* (A7) and *M. hmongorum* (A8). Admixture zones were widespread across central Indochina, the northern Annamites and Sumatra. Genetic diversity was concentrated at intermediate latitudes, declining toward both the southern (Peninsular Malaysia, Sumatra) and northern (Myanmar, northern Vietnam, Taiwan) edges of the distribution.

### Timetrees, discordances and historical gene flow

Timeframes of diversification varied among *Microhyla* clades, ranging from ca. 15 Ma in the *M. butleri* complex to <3 Ma in *M. pulchra* (Fig. 4). Divergence times estimated under the multispecies coalescent based on genomic SNPs were consistently younger than those inferred from mitochondrial datasets, as expected given that divergence at single loci – particularly mtDNA, owing to its smaller effective population size – typically predates species/lineage divergence and does not account for ILS among nuclear loci. The species trees were generally congruent with the ML reconstructions based on concatenated RAD loci including all samples, with discrepancies likely attributable to differences in methodological inferences and sampling schemes, particularly regarding the inclusion of admixed individuals, which are known to distort tree topology (Ambu et al. 2023).

**Fig. 4.**
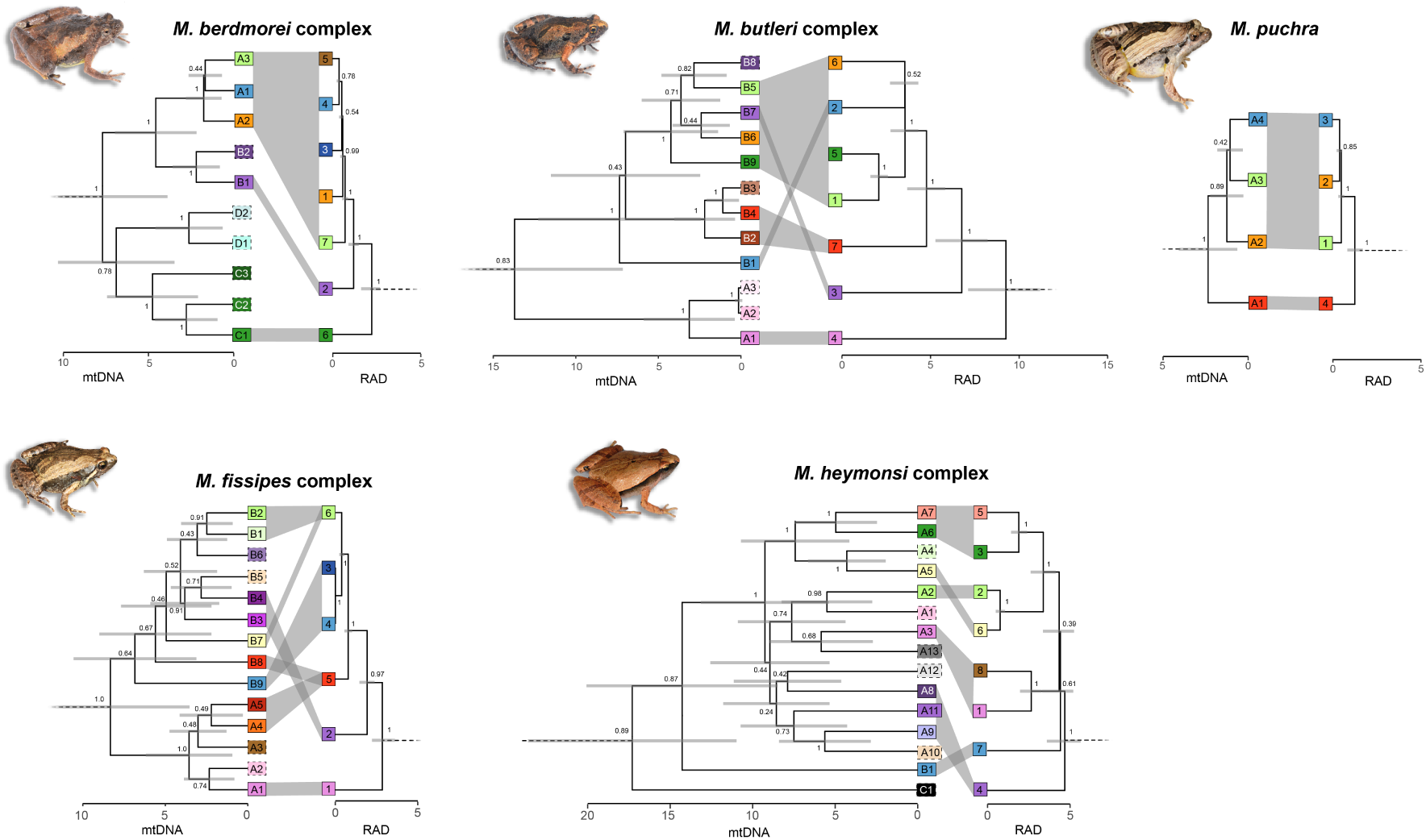
Mitochondrial and nuclear timetrees of *Microhyla* species/species complexes. Correspondence between lineages are shown by shaded areas. Dash frames indicate mtDNA lineages not sampled for ddRAD-seq. Lineage color and labels correspond to those in Figs. 1–2.

Nuclear and mitochondrial trees differed in the placement of several lineages. While some conflicting topologies may reflect insufficient phylogenetic resolution – especially in mtDNA datasets for recent divergences – deeper cyto-nuclear discordances more likely reflect historical hybridization involving mitochondrial capture or lineage fusion, especially when associated with strongly supported nodes. Historical gene flow was further suggested by significant excesses of shared derived alleles detected among several lineages in all clades by the ABBA–BABA analyses (*f_b_* up to 0.22; Fig. 5) and was often associated with cyto-nuclear discordances and weakly supported nodes in the species trees (Fig. 4).

**Fig. 5.**
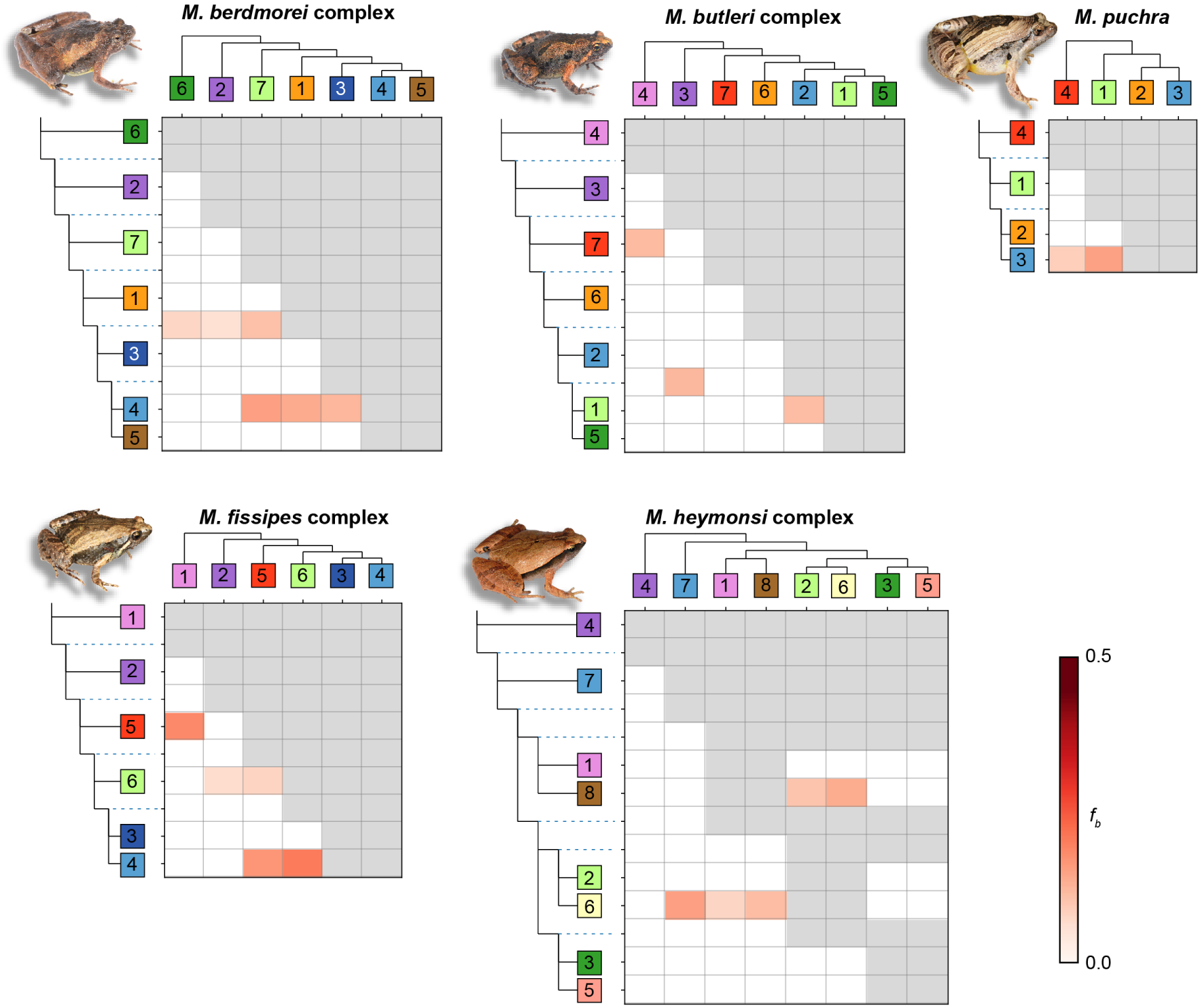
Heatmap of significant *f_b_* statistics in *Microhyla* species/species complexes. Backbone topologies are shown, with ancestor lineages as dash lines. Lineage color and labels correspond to those in Figs. 1–2.

The *M. butleri* complex provides a compelling example. In this clade, the placement of the Myanmar lineage within the Indochinese mitochondrial diversity (B7), despite its deep nuclear divergence (cluster 3), likely reflects mitochondrial capture during historical hybridization. Consistent with this scenario, the Myanmar lineage shares derived nuclear alleles with western and southwestern Thailand lineages (clusters 1 and 5). Reciprocally, the shallow nuclear divergence of the southern Vietnamese lineage (cluster 2), despite its deeply divergent mtDNA lineage (B1), suggests recent introgression from neighboring populations; the western Thailand lineage (cluster 1) accordingly shares a significant proportion of derived alleles with South Vietnam. Cyto-nuclear discordances and historical gene flow appear most prevalent in the *M. fissipes* complex, where nuclear genomes seem homogenized despite deep mitochondrial divergence, and distinct mitogroups segregate in single nuclear clusters (Figs. 4–5). In the *M. heymonsi* complex, at least two pairs of nuclear lineages show conflicting placements in the mtDNA tree (clusters 7/4 and 2/6; Fig. 4) and exhibit significant *f_b_* estimates (Fig. 5). In the *M. berdmorei* complex and *M. pulchra*, discrepancies between mtDNA and nuclear trees are restricted to recently diverged lineages (with weakly supported nodes) that are currently admixing in contact zones, and may therefore reflect either ILS or previous episodes of hybridization. Historical gene flow also affects deeper lineages without cyto-nuclear discordance in several complexes (Fig. 5).

## Discussion

### Shared biogeographic drivers of diversification

Using dense mitochondrial sampling combined with genome-scale analyses, our study reconstructed the comparative phylogeographic history of five co-distributed *Microhyla* species complexes throughout the Indochinese biodiversity hotspot. Despite having diverged from one another approximately 20–30 million years ago (Gorin et al. 2020), these lineages exhibit strikingly congruent geographic structuring. Across all groups, major lineage breaks repeatedly occur between Myanmar, the Malay Peninsula, southern Vietnam and northern Vietnam/southern China, while the geographically most widespread complexes further diversified into southeastern China/Taiwan and Sundaland. Such repeated phylogeographic breaks strongly suggest that common eco-geographical drivers have recurrently structured diversification across taxa, supporting the hypothesis that Indochina functions as a mosaic of biogeographic units whose boundaries have repeatedly promoted lineage persistence and divergence.

The mechanisms generating this mosaic remain difficult to disentangle, but our results are consistent with a hierarchical interplay between long-term geological processes and more recent climatic oscillations. At deeper timescales, the uplift of the Himalayan– Tibetan Plateau and the associated establishment and intensification of the Asian monsoon system fundamentally reorganized regional climates during the Miocene and Pliocene (Clift & Plumb 2008; Ao et al. 2021; Holbourn et al. 2021). Increasing rainfall seasonality likely reshaped the distribution of humid forests, seasonal forests and more open habitats across mainland Southeast Asia, creating persistent environmental gradients that may have promoted lineage divergence. Superimposed on this framework, repeated Plio-Pleistocene climatic oscillations further altered forest connectivity through cycles of contraction and expansion, modifying opportunities for isolation and dispersal across mainland Southeast Asia (Cannon et al. 2009; Hughes et al. 2011; Morley 2018; Raes et al. 2014). Mountain systems such as the Annamites, Kayah–Karen, Tenasserim and Indo-Burman ranges may have acted as long-term climatic refugia maintaining humid environments during periods of increased aridity, whereas intervening lowland basins and river systems, including the Irrawaddy, Chao Phraya and Mekong drainages, probably alternated between barriers and corridors as environmental conditions changed. The widespread admixture zones recovered by our genomic analyses are consistent with repeated episodes of secondary contact following these cycles of isolation and reconnection.

Despite these shared phylogeographic breaks, the five *Microhyla* complexes differ markedly in the depth and complexity of their diversification. Whereas *M. pulchra* and *M. fissipes* exhibit only shallow phylogeographic structure (notably in the nuclear DNA), other complexes comprise numerous deeply divergent lineages. These differences imply that species have experienced distinct biogeographic histories within the same regional landscape. The young and weakly-diversified complexes may have established their current range only recently, and from one or a few historical refugia, leaving insufficient time for populations to accumulate deep divergence, whereas the older and more structured complexes probably experienced long-term persistence and diversification across multiple refugia. These alternative histories could be further modulated by differences in ecological specialization and dispersal ability. Although the ecology of the five *Microhyla* complexes remains poorly documented, their widespread sympatry across much of Indochina raises the possibility of ecological differentiation, including microhabitat partitioning and climatic niche shifts. Comparative species distribution modelling coupled with paleodistribution reconstructions represents a promising avenue to disentangle the respective contributions of historical range dynamics and ecological traits to diversification.

### Hybridization as a hallmark of dynamic biogeographic history

While phylogeographic studies of Palearctic amphibians have extensively documented historical introgression and reticulation (e.g. Ambu et al. 2025; Dufresnes et al. 2019; Mars et al. 2025; Koster et al. 2026), the contribution of hybridization to the evolution of tropical Southeast Asian amphibians has only recently begun to receive attention (Chan et al. 2020). Amphibian diversification in the region is commonly portrayed as resulting from long-term allopatric persistence within environmentally stable refugia, giving rise to the numerous regional endemic and microendemic species that characterize the Indochinese herpetofauna (reviewed by Poyarkov et al. 2021, 2023). Our phylogenomic analyses instead reveal a more dynamic evolutionary history. All five *Microhyla* complexes exhibit signatures of historical gene flow, further reflected by pronounced cytonuclear discordance in several groups (*M. butleri*, *M. fissipes* and *M. heymonsi*). Rather than representing isolated lineages evolving independently “on the spot”, populations appear to have repeatedly come into secondary contact and exchanged genes following periods of divergence. Beyond accumulating introgressed variation, the remarkable genomic plasticity of *Microhyla* is further illustrated by the recent discovery of an autotetraploid lineage of *M. fissipes* on Hainan Island arising through genome duplication (Chen et al. 2025). Although comparable phylogenomic evidence remains scarce for Southeast Asian amphibians, historical introgression has now been demonstrated in the ranid *Pulchrana picturata* complex (Chan et al. 2020), the bufonid *Duttaphrynus melanostictus* complex (Dufresnes et al. 2025), as well as in other tropical vertebrates, notably gibbons (*Hylobates*) (Matsudaira et al. 2021) and macaques (*Macaca*) (Fan et al. 2018). Historical introgression may therefore prove to be as pervasive in tropical as in temperate amphibians as genome-scale phylogenomic studies explicitly modelling gene flow progressively replace traditional phylogeographic analyses based on a handful of loci. Rather than a static assemblage of phylogeographic breaks, the Indochinese biodiversity hotspot should be viewed as a dynamic biogeographic system in which repeated cycles of isolation, secondary contact and introgression have continually reshaped lineage boundaries and generated biodiversity.

### Towards phylogeography-based taxonomy and conservation of the Indochinese diversity hotspot

Another pervasive feature of the Asian herpetofauna, amply illustrated here, is the occurrence of deeply divergent cryptic lineages within many traditionally recognized species (Poyarkov et al. 2020, 2022; Xu et al. 2024). This renders current taxonomy increasingly inadequate and hampers conservation, as unnamed evolutionary lineages are generally overlooked by legislation and management frameworks (Garnett & Christidis 2017; Pollock et al. 2020; Liu et al. 2022). At the same time, taxonomic revision of lineage-rich complexes is particularly challenging because species delimitation, specimen comparisons, and the allocation of available names all require a robust understanding of lineage relationships, distributions, and evolutionary independence. Although mitochondrial barcoding has proved invaluable for revealing hidden diversity, its inherent limitations – including incomplete lineage sorting, introgression and mitochondrial capture – can bias taxonomic inference, particularly in amphibians (Dufresnes & Jablonski 2022). Genome-scale validation of taxonomic hypotheses is therefore becoming essential (Vences et al. 2024), yet remains uncommon (Miralles et al. 2024). Importantly, genomic data not only facilitate the recognition of overlooked species, but also help avoid taxonomic inflation by identifying phylogeographic lineages that remain connected through substantial gene flow and therefore represent intraspecific diversity (Dufresnes et al. 2023; Gippner et al. 2026).

In this respect, additional genomic sampling in the *M. berdmorei* complex corroborates the recent recognition of four species (Trofimets et al. 2024), with the geographically extensive *M. malcolmi* comprising widely admixing, therefore conspecific phylogeographic lineages. Distinct candidate species are also found within the *M. butleri* and *M. heymonsi* complexes. Particularly striking is the Myanmar lineage of *M. butleri*, whose complete mitochondrial replacement masks a deeply divergent nuclear lineage, making it a likely example of a “super-cryptic species” sensu Dufresnes et al. (2019). In the *M. heymonsi* complex, phylogenomics supported deep nuclear lineages with absent or restricted introgression, namely *M. daklakensis* micro-endemic from south Vietnam, eastern Indochinese – southeastern Chinese populations (including Taiwan), northeastern Indochinese populations, and Peninsular Malaysian – Sundaland populations. Conversely, several recently described taxa appear to represent geographically restricted mitochondrial lineages nested within these broader candidate species. For instance, *M. nakkavaram* from the Andaman Islands and Indonesia belong to a Peninsular Malaysian–Sundaland nuclear phylogroup, whereas *M. ninhthuanensis* corresponds to a unique southern Vietnamese mitochondrial lineage that broadly admix with other south Indochinese lineages. Likewise, in the *M. fissipes* complex, the deep mitochondrial split between southeastern Chinese (*M. fissipes*) and Indochinese (*M. mukhlesuri*) populations contrasts with much shallower nuclear divergence and a seemingly broad admixture zone in northern Vietnam. Pending genomic assessment of mainland Chinese populations, these lineages may be more appropriately regarded as components of a single species (e.g., subspecies). The *Microhyla* diversifications thus highlight how reliance on mitochondrial data alone may simultaneously underestimate species diversity by overlooking deeply divergent nuclear lineages and overestimate it by assigning species status to geographically structured mitochondrial haplogroups. Comparative phylogenomics offers a much more robust framework for distinguishing independently evolving species from structured populations within species, thereby improving both taxonomic stability and the evolutionary relevance of species classifications.

Finally, the partially shared spatial distribution of genetic diversity across *Microhyla* complexes identifies regions that deserve particular conservation attention as hotspots of evolutionary potential. Whereas conservation priorities have traditionally focused on species richness and endemism, areas concentrating exceptional levels of genetic diversity should also receive attention as genetic diversity underpins population, and therefore species and ecosystem resilience (Shaw et al. 2026). The southern Annamite Mountains emerge as the strongest candidate, harboring the highest genomic diversity in four of the five complexes investigated. Likewise, the Tenasserim Hills represent an additional hotspot for *M. pulchra* and the *M. butleri* and *M. heymonsi* complexes. Besides supporting numerous endemic lineages, these regions appear to function as long-term evolutionary reservoirs in which populations have persisted, diverged and repeatedly reconnected through time. Conserving such centers of genetic variation shall thus help preserve the evolutionary processes and adaptive potential that generate and maintain biodiversity, in an effort to reverse the biodiversity crisis that is particularly acute in Southeast Asia (Struebig et al. 2025).

## Acknowledgements

The authors are grateful to T Ruangsuwan, T Worranuch (Thailand), T Matsukoji (Japan), B Zhang, D Zou, K Jiang, F Yan, L-J Wang, H Zhao, and N Zhu for their help during the field surveys; to the Animal Bank at the Germplasm Bank of Wild Species (https://cstr.cn/31121.02.GBOWS) for providing biological materials and technical support; to Amy Lathrop (Royal Ontario Museum, Canada) for the provision of tissue grants; to members of MSU HerpLab, including SS Idiiatullina and NS Kliukin, for their support and assistance; to the Laboratory Animal Research Center, University of Phayao, and the Institute of Animal for Scientific Purposes Development (IAD), Thailand, for permission to do fieldwork in Thailand; and to the local forestry departments in China for granting permissions for field surveys and specimen collection, as well as for their field assistance. NAP is grateful to AN Kuznetsov, SP Kuznetsova, HD Nguyen, and LP Korzoun for their support and organization of fieldwork. Specimen collection and animal use protocols were approved by the Institutional Ethical Committee of Animal Experimentation of the University of Phayao, Phayao, Thailand (certificate number UP-AE64-02-04-005; to CS), and were strictly compliant with the ethical conditions of the Thailand Animal Welfare Act. Fieldwork, including the collection of animals in the field, was authorized by the Institute of Animals for Scientific Purpose Development (IAD), Bangkok, Thailand (n°U1-01205-2558 and UP-AE59-01-04-0022; to C. Suwannapoom). Permission to conduct fieldwork in Vietnam was granted by the Bureau of Forestry, the Ministry of Agriculture and Rural Development of Vietnam, and local administrations. Please see Supplementary Table S1 for details on all permits. The fieldwork was completed within the framework of, and with partial financial support from, the research project “Conservation, restoration, and sustainable use of tropical forest ecosystems based on the study of their structural and functional organization” of the Joint Vietnam-Russia Tropical Science and Technology Research Center and was partially supported by this project during 2009-2023. This work was supported by the Russian Science Foundation (grant n°22-14-00037-P to NAP; sampling, data analyses); the Unit of Excellence 2026 (Grant No. UoEICA002; to CS); the Thailand Science Research and Innovation Fund and the University of Phayao (Unit of Excellence 2027 on Aquatic animals biodiversity assessment [Phase III] to CS; sampling) and the National Natural Science Foundation of China (Research Fund for International Scientists n°3211101356 to CD; labwork), the Key R & D program of Yunnan Province, China (n°202503AP140029 to JC; sampling and labwork), the Yunnan Revitalization Talent Support Program Yunling Scholar (to JC; sampling and labwork), and China’s Biodiversity Observation Network (Sino-BON; to JC; sampling and labwork).

